# Swarm learning for decentralized artificial intelligence in cancer histopathology

**DOI:** 10.1101/2021.11.19.469139

**Authors:** Oliver Lester Saldanha, Philip Quirke, Nicholas P. West, Jacqueline A. James, Maurice B. Loughrey, Heike I. Grabsch, Manuel Salto-Tellez, Elizabeth Alwers, Didem Cifci, Narmin Ghaffari Laleh, Tobias Seibel, Richard Gray, Gordon G. A. Hutchins, Hermann Brenner, Tanwei Yuan, Titus J. Brinker, Jenny Chang-Claude, Firas Khader, Andreas Schuppert, Tom Luedde, Sebastian Foersch, Hannah Sophie Muti, Christian Trautwein, Michael Hoffmeister, Daniel Truhn, Jakob Nikolas Kather

## Abstract

Artificial Intelligence (AI) can extract clinically actionable information from medical image data. In cancer histopathology, AI can be used to predict the presence of molecular alterations directly from routine histopathology slides. However, training robust AI systems requires large datasets whose collection faces practical, ethical and legal obstacles. These obstacles could be overcome with swarm learning (SL) where partners jointly train AI models, while avoiding data transfer and monopolistic data governance. Here, for the first time, we demonstrate the successful use of SL in large, multicentric datasets of gigapixel histopathology images comprising over 5000 patients. We show that AI models trained using Swarm Learning can predict *BRAF* mutational status and microsatellite instability (MSI) directly from hematoxylin and eosin (H&E)-stained pathology slides of colorectal cancer (CRC). We trained AI models on three patient cohorts from Northern Ireland, Germany and the United States of America and validated the prediction performance in two independent datasets from the United Kingdom using SL-based AI models. Our data show that SL enables us to train AI models which outperform most locally trained models and perform on par with models which are centrally trained on the merged datasets. In addition, we show that SL-based AI models are data efficient and maintain a robust performance even if only subsets of local datasets are used for training. In the future, SL can be used to train distributed AI models for any histopathology image analysis tasks, overcoming the need for data transfer and without requiring institutions to give up control of the final AI model.

## Introduction

Artificial intelligence (AI) is expected to have a profound impact on the practice of medicine in the next ten years.^1–4^ In particular, medical imaging is already in the process of being transformed by the application of AI solutions.^5^ Such AI solutions can automate manual tasks in medical image analysis, but can also be used to extract hidden information beyond what is visible to the human eye.^6,7^ In particular, digitized histopathology images contain a wealth of clinically relevant information which AI can extract.^3^ For example, deep convolutional neural networks have been used to predict molecular alterations of cancer directly from routine pathology slides.^8–13^ In 2018, a landmark study showed a first proof-of-principle for this technology in lung cancer.^8^ Since then, dozens of studies have extended and validated these findings to multiple tumor types including colorectal cancer (CRC)^9,14,15^, gastric cancer^16^, bladder cancer^10^, breast cancer^13^, among other tumor types^10–12,17,18^. These methods expand the utility of hematoxylin and eosin (H&E) stained tissue slides from routine tumor diagnosis and subtyping to a source for direct prediction of molecular alterations.^3^

AI models are data hungry. In particular, in histopathology, the performance of AI models in increases with the size and diversity of the training set.^16,19,20^ Training clinically useful AI models usually requires sharing of patient-related data with a central repository.^21,22^ In practice, such data sharing - especially across different countries - faces legal and logistic obstacles. Data sharing between institutions may require patients to forfeit their rights of data control. This problem has been tackled by (centralized) federated learning (FL)^23,24^, in which multiple AI models are trained independently on separate computers (peers). In FL, peers do not share any input data with each other and only share the learned model weights. However, a central coordinator governs the learning progress based on all trained models, monopolizing control and commercial exploitation.

In the last two years, this limitation of FL has been addressed by a new group of decentralized learning technologies, including blockchain FL^25^ and swarm learning (SL).^26^ In SL, AI models are trained locally and models are combined centrally without requiring central coordination. By using blockchain-based coordination between peers, SL removes the centralization in FL and raises all contributors to the same level. In the context of healthcare data analysis, SL leads to equality in training multicentric AI models and creates strong incentives to collaborate without concentrating data or models in one place. This could potentially facilitate collaboration between multiple parties, hence generating more powerful and more reliable AI systems. Ultimately, SL could improve the quality, robustness and resilience of AI in healthcare. However, SL has not been systematically applied to medical image data in oncology. In particular, it has not been applied to histopathology images, a common data modality with a high information density.^3^

To address this, we performed a retrospective multicentric study of SL for AI-based prediction of molecular alterations directly from conventional histology images. As pathology services are currently undergoing a digital transformation, embedding these AI methods into routine diagnostic workflows could ultimately enable pre-screening of patients, thereby reducing the number of costly genetic tests and increasing the speed by which results are available to clinicians.^27^ The prediction performance of such systems increases markedly by training on thousands rather than just hundreds of patients.^19,20^ We hypothesized that SL could be a substitute for centralized collection of large patient cohorts in histopathology, improving prediction performance^20^ and generalizability^22^ without centralizing data collection or control over the final model.

## Materials and Methods

### Ethics statement

This study was carried out in accordance with the Declaration of Helsinki. This study is a retrospective analysis of digital images of anonymized archival tissue samples from five cohorts of CRC patients. Collection and anonymization of patients in all cohorts took place in each contributing center. Ethical approval for research use of all cohorts was obtained from each contributing center.

### Patient cohorts

We collected digital whole slide images (WSI) of H&E-stained slides of archival tissue sections of human colorectal cancer (CRC) from five patient cohorts (clinico-pathological characteristics in **Table 1**). First, the Northern Ireland Epi700 (n=661, **Suppl. Figure S1**) cohort study on colon cancer of only patients with stage II-III cancer, provided by the Northern Ireland Biobank^28,29^ (application NIB20-0346). Second, the “Darmkrebs: Chancen der Verhütung durch Screening” study (DACHS, n=2448, **Suppl. Figure S2**), a large population-based case-control study, including samples of CRC patients at different disease stages recruited at >20 hospitals in Germany, which is coordinated by the German Cancer Research Center (DKFZ, Heidelberg, Germany)^30–32^. Third, “The Cancer Genome Atlas” (TCGA) CRC cohort (n=632, **Suppl. Figure S3**), a large collection of tissue specimens from multiple study centers across different countries, but largely from the United States of America (USA).^33^ Fourth, the “Quick and Simple and Reliable” (QUASAR) trial (n=2206, **Suppl. Figure S4**), which originally aimed to determine survival benefit from adjuvant chemotherapy in CRC patients from the United Kingdom (UK)^34^. Fifth, the Yorkshire Cancer Research Bowel Cancer Improvement Programme^35^ (YCR-BCIP) cohort (n=889 surgical resection slides, **Suppl. Figure S5**), collected in the Yorkshire Region in the UK. For all cohorts, *BRAF* mutational status and microsatellite instability (MSI) / mismatch repair deficiency (dMMR)^36^ data were acquired. In YCR-BCIP, analysis of *BRAF* was only undertaken for dMMR tumors and *BRAF* mutational status was therefore not assessed in this cohort in the current study.

**Table 1:** Clinico-pathological features of all cohorts. Number of patients (N Patients), Interquartile range (IQR), microsatellite instability (MSI), microsatellite stability (MSS), b-Raf kinase mutational status (BRAF), wild type (wt), mutated (mut), not available (N/A). Right-sided CRC from cecum to transverse colon. * BRAF testing in YCR-BCIP was only performed in MSI/dMMR cases and was therefore not used as a prediction target in this study.

|  | TCGA | DACHS | Epi700 | YCR-BCIP | QUASAR |
| --- | --- | --- | --- | --- | --- |
| N Patients | 632 | 2448 | 661 | 889 | 2190 |
| Age (median) | 68 | 69 | 72.7 | 71 | 63 |
| Age (IQR) | 18 | 14 | 14.5 | 15 | 12 |
| Gender: Male | 322 (50.9%) | 1436 (58.7%) | 358 (54.2%) | 494 (55.6%) | 1334 (60.9%) |
| Gender: Female | 292 (46.2%) | 1012 (41.3%) | 303 (45.8%) | 395 (44.4%) | 848 (38.7%) |
| Gender: Unknown | 18 (2.85%) | 0 | 0 | 0 | 8 (0.4%) |
| MSS/pMMR | 392 (62%) | 1836 (75%) | 471 (71.3%) | 760 (85.5%) | 1529 (69.8%) |
| MSI/dMMR | 65 (10.3%) | 210 (8.6%) | 136 (20.6%) | 129 (14.5%) | 246 (11.2%) |
| unknown MS/MMR status | 175 (27.7%) | 402 (16.4%) | 54 (8.1%) | 0 | 415 (19%) |
| MSI/MMR method | PCR 5-plex | PCR 3-plex | PCR | IHC | IHC |
| BRAF status: wt | 471 (74.5%) | 1930 (78.8%) | 553 (83.7%) | 32 (3.6%,*) | 1358 (62%) |
| BRAF status: mut | 63 (10%) | 151 (6.2%) | 92 (13.9%) | 75 (8.4%,*) | 129 (5.9%) |
| BRAF status: unknown | 98 (15.5%) | 367 (15%) | 16 (2.4%) | 782 (88%) | 916 (41.8%) |
| Stage 1 | 76 (12%) | 485 (19.8%) | 0 | 169 (19%) | 5 (0.2%) |
| Stage 2 | 166 (26.3%) | 801 (32.7%) | 394 (59.6%) | 317 (35.7%) | 53 (2.4%) |
| Stage 3 | 140 (22.2%) | 822 (33.6%) | 267 (40.4%) | 370 (41.6%) | 1653 (75.5%) |
| Stage 4 | 63 (10%) | 337 (13.8%) | 0 (0%) | 33 (3.7%) | 268 (12.2%) |
| Stage unknown | 187 (29.5) | 3 (0.1%) | 0 | 0 | 211 (9.7%) |
| Left-sided CRC | 248 (39.2%) | 1607 (65.6%) | 280 (42.3%) | 487 (54.8%) | 1158 (52.9%) |
| Right-sided CRC | 176 (27.8%) | 819 (33.5%) | 375 (56.7%) | 332 (37.3%) | 754 (34.4%) |
| Unknown side | 209 (33%) | 22 (0.9%) | 6 (1%) | 70 (7.9%) | 278 (12.7%) |

### Deep learning and swarm learning method

For prediction of molecular features from image data, we adapted our weakly-supervised prediction pipeline “Histology Image Analysis (HIA)”^9^ which was demonstrated to outperform similar approaches for mutation prediction in a recent benchmark study.^37^ Briefly, the workflow entails the following steps: As preprocessing step, high resolution WSIs were tessellated into patches of size (512 × 512 × 3) pixels and color-normalized.^38^ During this process, blurry patches and patches with no tissue are removed from the data set using canny edge detection in OpenCV.^37^ Subsequently, we used ResNet18 to extract a (512 × 1) feature vector from 150 randomly selected patches for each patient, as previous work showed that 150 patches are sufficient to obtain robust predictions.^9^ Feature vectors and patient-wise target labels (BRAF or MSI status) served as input to a fully connected classification network (FCN). The FCN comprised four layers with (512×256), (256×256), (256×128) and (128×2) connections with a ReLU activation function (hyperparameters in **Suppl. Table S1**). The swarm learning architecture had four components, running as separate processes (nodes) on separate bare-metal servers (peers) in a Docker container. The key component is the Swarm Learning (SL) process, containing HIA (**Suppl. Figure S6A**). During training in the SL process on three devices, the network weights are sent to a Swarm Network (SN) process, which coordinates peer crosstalk and sends back averaged weights to the SL process. Peer crosstalk happens once every synchronization interval (“sync interval”). Averaging and exchange of weights is coordinated by an Ethereum blockchain.^39^ For identity management, a Secure Production Identity Framework for Everyone (SPIFFE) running on its own virtual environment (SPIFFE Runtime Environment, SPIRE) serves as the third process, generating a secured gateway between peers. Formal hyperparameter optimization was performed for the sync interval for the task of MSI status prediction (**Suppl. Figure S6D-E**). Technical details are available in **Suppl. Methods**.

### Experimental Design and Statistics

Throughout the study, TCGA, Epi700, DACHS were used as training cohorts and QUASAR and YCR-BCIP were used as external test cohorts (**Table 1**). First, we trained MSI and *BRAF* classifiers on each of the training cohorts individually. Second, all training cohorts were merged and new classifiers were trained on the merged cohort. Third, classifiers were trained by SL, with the SL training process being initiated on three separate bare metal servers containing one training cohort each. Finally, all models were externally validated on the validation cohorts. Two variants of SL were explored: baseline SL and weighted SL. For baseline SL, each cohort was trained for a fixed number of epochs and two resulting models were saved at two checkpoints, b-chkpt1 and b-chkpt2. B-chkpt 1 was reached when the smallest cohort concluded the final epoch. B-chkpt2 was reached when the second-smallest cohort concluded the final epoch. In baseline SL, all cohorts were assigned an equal weight. This is motivated by the fact that at the start of training, partners might not know the total amount of data they contribute because more data can be dynamically added during training. On the other hand, if partners know exactly how much data they will contribute, differences in cohort sizes can be compensated by applying proportional weights. Hence, in weighted SL, smaller cohorts are trained for more epochs than larger cohorts, but contribute proportionally less to the final model. In weighted SL, only one model checkpoint is generated, w-chkpt. Finally, to investigate data efficiency, we repeated all experiments for subsets of 25, 50, 100, 200, 300 and 400 patients per cohort, randomly selected in a stratified way (preserving class proportions). All experiments were repeated five times with different random seeds. The primary endpoint for this study was the area under the receiver operator characteristic curve (AUROC) for detection of binary categorical outputs. The AUROCs of five training runs of a given model were compared. A two-tailed unpaired t-test with p<= 0.05 was considered statistically significant. In the manuscript, AUROCs are given as mean +/- standard deviation. All raw results of all experimental repetitions are available in **Suppl. Table S2**.

### Code availability

All source codes for the baseline histology image analysis (HIA) workflow are available at https://github.com/KatherLab/HIA. Source codes for Swarm-HIA are available at https://github.com/KatherLab/SWARM. We built our codes on top of the “SL community edition” by Hewlett Packard Enterprise (HPE, Spring, Texas, United States), which is publicly available under an Apache 2.0 license at https://github.com/HewlettPackard/swarm-learning.

## Results

### Histology image analysis workflows can be coupled with swarm learning

In this study, we aimed to develop a swarm-learning-capable histopathology AI model for molecular classification of solid tumors based on histopathology images (**Figure 1A, Suppl. Figure S6A**). We collected three large datasets (**Figure 1B, Figure 2A**) in three physically separate computing servers (**Suppl. Figure S6B**), integrated an end-to-end histopathology AI pipeline with a swarm callback and optimized the synchronization (sync) interval between peers, i.e. different physical computers (**Suppl. Figure S6C**). We found that synchronizing learning progress between peers once every four iterations (**Figure 2B-C**) yielded the highest performance in a benchmark task (MSI prediction, **Suppl. Figure S6D**) while retaining a low overall training time (**Suppl. Figure S6E**). Total training time was inversely proportional to the sync interval and was 01:17:41(hours:minutes:seconds) for a sync interval of 1, compared to 00:18:40 and 00:09:26 for sync intervals of 4 and 8 iterations, respectively, indicating that the swarm learning time was dominated by network communication overhead (**Suppl. Figure S6E**).

**Figure 1:**
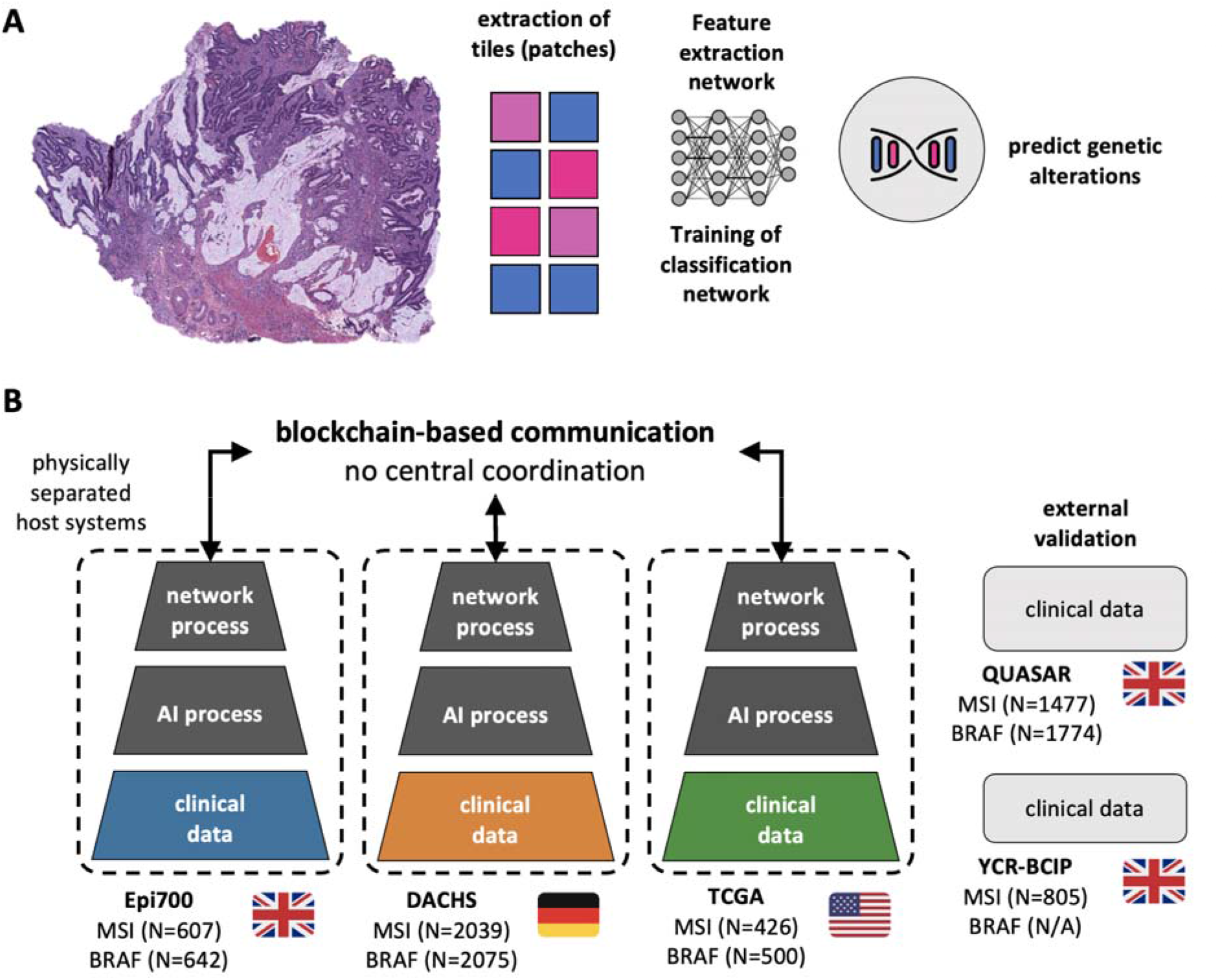
Schematic of the swarm learning workflow and experimental layout. **(A)** Histology image analysis (HIA) workflow, **(B)** swarm learning workflow and cohorts included in this study. On three physically separate bare-metal servers (dashed line), three different sets of clinical data reside. Each server runs an AI process (a program that trains a model on the data) and a network process (a program that handles communication with peers via blockchain). Icon sources: openmoji, Twitter Twemoji (CC-BY).

**Figure 2:**
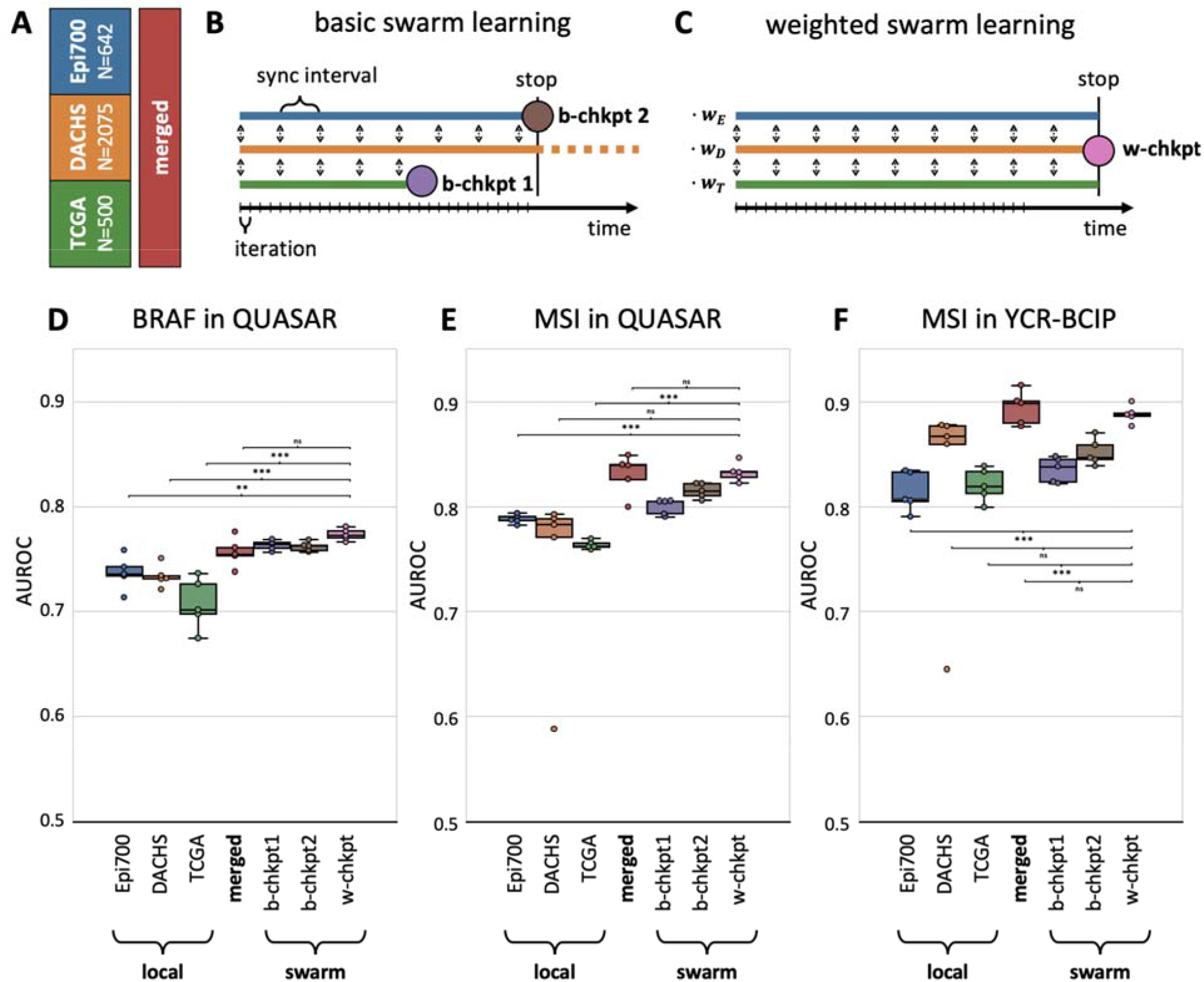
Model performance for local models vs. swarm models for *BRAF* prediction. **(B)** Schematic of the basic swarm learning (SL) experiment and the **(C)** weighted SL experiment. **(D)** Classification performance (area under the receiver operating curve, AUROC) for prediction of *BRAF* mutational status on a patient level in the QUASAR data set, displayed as box plot (box shows the median and quartiles as the whiskers expand to the rest of the distribution, with the exception of points identified as outliers) as well as all original data points. **(E)** AUROC for prediction of MSI status in QUASAR and **(F)** prediction of MSI status in the YCR-BCIP dataset. Abbreviations: chkpt = checkpoint, chkpt 1, 2 = swarm checkpoint 1 and 2, respectively. W-chkpt = Total cohort sizes (N patients) were: 642 for Epi700, 2075 for DACHS, 500 for TCGA (*BRAF*) and 0.0014 for Epi700, 8.65E-05 for DACHS and 0.0003 for TCGA (MSI). *: p<0.05, **: p<0.01, ***: p<0.001, ns: p>0.05.

### Swarm learning performance is on par with models trained on the merged dataset

We then used our swarm-capable histology image analysis model in a retrospective multicenter study for prediction of *BRAF* mutational status from colorectal cancer (CRC) histopathology WSIs. First, we trained local AI models on each of three training cohorts separately: Epi700 (N=594 patients from Northern Ireland), DACHS (N=2039 patients from southwest Germany) and TCGA (N=426, **Figure 1B**). Evaluating the patient-level prediction performance of these local models on the QUASAR cohort (N=1774 patients from the UK), we found that these models achieved AUROCs of 0.7358 (+/- 0.0162), 0.7339 (+/- 0.0107) and 0.7071 (+/- 0.0243) when trained on Epi700, DACHS and TCGA alone, respectively (**Figure 2D**). Merging the three training cohorts on a central server (centralized model) significantly improved prediction AUROC to 0.7567 (+/- 0.0139, p=0.0727 vs. Epi700, p=0.0198 vs. DACHS, p=0.0043 vs. TCGA**, Suppl. Table S3**). This was compared to the performance of SL-AI models: B-chkpt1 was obtained when the partner with the smallest training cohort (TCGA) reached the last epoch (**Figure 2B**). This model achieved a prediction AUROC on the test set of 0.7634 (+/- 0.0047), which was significantly better than each local model (p=0.0082 vs. Epi700, p=0.0005 vs. DACHS, p=0.0009 vs. TCGA), but not significantly different from the merged model (p=0.3433, **Figure 2D**). B-chkpt2, which was obtained when the partner with the second-smallest training cohort reached the last epoch, achieved a similar performance: This model achieved an AUROC of 0.7621 (+/- 0.0045), which was significantly better than each local model (p=0.0105 vs. Epi700, p=0.0006 vs. DACHS, p=0.0011 vs. TCGA), and on par with the merged model (p=0.4393, **Figure 2D**). We validated these findings in a different task, prediction of MSI/dMMR status in the QUASAR (**Figure 2E**) and YCR-BCIP (**Figure 2F**) cohort. In QUASAR, b-chkpt1 and b-chkpt2 achieved prediction AUROCs of 0.8001 (+/- 0.0073) and 0.8151 (+/- 0.0071), respectively and thereby significantly outperformed single-cohort models trained on Epi700 with an AUROC of 0.7884 (+/-0.0043 (p=0.0154 and p=8.79E-05 for B-chkpt1 and −2, respectively, **Suppl. Table S4**). Similarly, SL outperformed MSI prediction models trained on TCGA with an AUROC of 0.7639 (+/-0.0162) (p=1.09E-05 and p=6.14E-07 for b-chkpt1 and2, respectively). However, there was no significant difference between the model trained on the largest data set DACHS compared to b-chkpt1 or −2 in QUASAR (**Figure 2E**) and YCR-BCIP (**Figure 2F**). As another variation of SL, we added adjustable cohort weights proportional to the cohort size and obtained the weighted SL-AI model w-chkpt. For prediction of *BRAF* mutational status, w-chkpt achieved an AUROC of 0.7736 (+/- 0.0057). This represented a significant improvement compared to all other models, including the local models of Epi700 (p=0.0015), DACHS (p=8.65E-05), TCGA (p=0.0004), but also the merged model (p=0.0374) and b-chkpt1 (p=0.0154) and b-chkpt2 (p=0.0081, **Figure 2D, Suppl. Table S3**). For MSI prediction in QUASAR, w-chkpt significantly outperformed the local Epi700 model (p=8.93E-06) and the local TCGA model (p=2.83E-07) while the performance differences compared to the DACHS model were not statistically significant (DACHS AUROC 0.8326 [+/- 0.0090] vs. w-chkpt AUROC 0.7403 [+/- 0.0878], p=0.05705, **Figure 2E, Suppl. Table S4**). Similar results were obtained for the second MSI validation dataset (**Figure 2F, Suppl. Table S5**). Compared to the merged model, w-chkpt was not significantly different for MSI prediction in QUASAR (merged AUROC 0.8308 [+/- 0.0190] vs. w-chkpt AUROC 0.8326 [+/- 0.0089], p=0.8650) or MSI prediction in YCR-BCIP (merged AUROC 0.8943 [+/- 0.0161] vs. w-chkpt AUROC 0.8882 [+/- 0.0084], p=0.4647), i.e. the merged model and w-chkpt performed on par (**Figure 2E-F**). Together, these data show that swarm-trained models consistently outperform local models and perform on par with centralized models in pathology image analysis.

### Swarm learning models are data-efficient

Learning from small datasets is a challenge in medical AI because prediction performance generally increases with increasing size of the training dataset.^19,20^ Therefore, we investigated whether SL could compensate for the performance loss which occurs when only a small subset of patients from each institution is used for training. We found that restricting the patient number in each training set to 400, 300, 200 and 100 markedly reduced prediction performance for single-dataset models. For example, for prediction of *BRAF* mutational status in QUASAR, training on only a subset of patients in Epi700, DACHS or TCGA markedly reduced prediction performance and increased the model instability as measured by interquartile range (IQR) of predictions in experimental repetitions (**Figure 3A, Suppl. Table S6**). In particular, for training *BRAF* prediction models on the largest cohort DACHS there was a pronounced performance drop in AUROC from training on all patients (AUROC 0.7339 [+/- 0.0108]) to an AUROC of 0.6626 [+/- 0.0162] when restricting the number of patients in the training set to 200 patients. Performance losses for the model which trained on the centrally merged data were less pronounced down to a number of 50 patients per cohort (**Figure 3A**). Strikingly, swarm learning was also able to rescue the performance: down to 100 patients per cohort, weighted swarm learning (w-chkpt) maintained a high performance of 0.7000 (+/- 0.0260) for 100, 0.7139 (+/- 0.0149) for 200 and 0.7438 (+/- 0.0093) for 300 patients. These models were not statistically significantly different from the merged model (p=0.7726, p=0.7780, p=0.2719, p=0.7130 for 100, 200, 300, 400 patients, respectively, **Figure 3A**). Similarly, b-chkpt1 and b-chkpt2 maintained a high performance (comparable to the merged model) down to 100 patients per cohort. For MSI prediction in QUASAR, w-chkpt was comparable to the merged model down to 300 patients per cohort (p=0.4342 and p=0.7847 for 300 and 400 patients, respectively). For 200 patients or less, the merged model outperformed local models and swarm models (**Figure 3B, Suppl. Table S7**). Similarly, for MSI prediction in YCR-BCIP, single-cohort performance dropped as patients were dropped from the training set but the merged models and swarm models could partially rescue this performance loss, although the merged model outperformed the swarm in this experiment(**Figure 3C, Suppl. Table S8**). All raw data are available in **Suppl. Table S2**. Together, these data show that swarm learning models are highly resilient to small training datasets for prediction of *BRAF* mutational status and partially resilient to small training datasets for prediction of MSI status.

**Figure 3:**
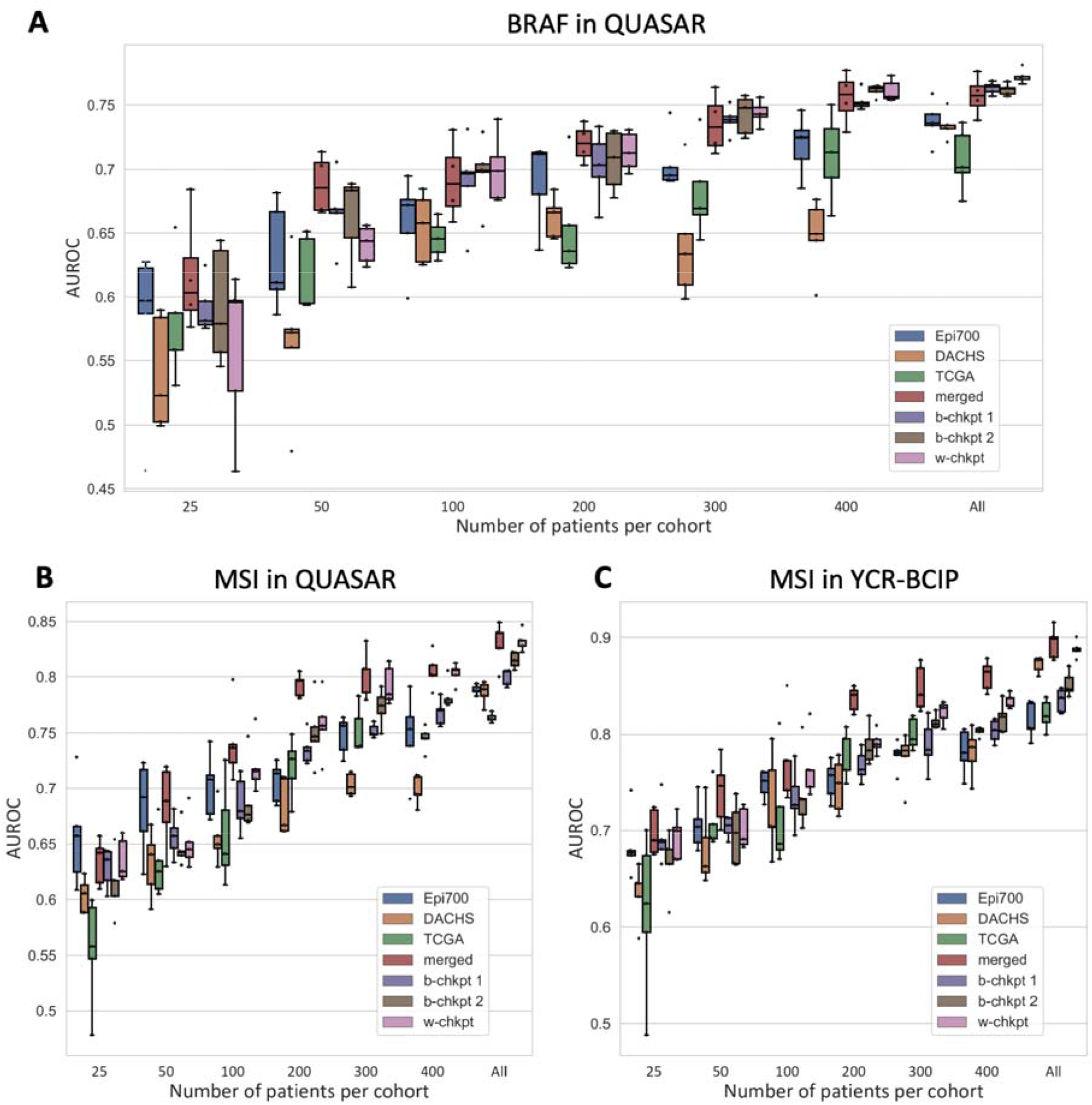
Model performance for local models vs. swarm models for microsatellite instability (MSI) / mismatch repair deficiency (dMMR) prediction. **(A)** Classification performance (area under the receiver operating curve, AUROC) for prediction of MSI mutational status on a patient level in the QUASAR cohort. **(B)** Classification performance (area under the receiver operating curve, AUROC) for prediction of dMMR mutational status on a patient level in the YCR-BCIP cohort. N patients per cohort (shown as “All” on the horizontal axis): 604 for Epi700, 2039 for DACHS, 426 for TCGA.

### Swarm learning models are interpretable and learn plausible patterns

Medical AI models should not just give a high performance but should also be interpretable.^40^ We investigated explainability by extracting the highest-scoring image patches for models trained on all individual training cohorts (**Figure 4A-C**), the merged cohort (**Figure 4D**) and the swarm models b-chkpt1 and b-chkpt2 (**Figure 4E-F**). In most cases there was a histological phenotype known to be associated with *BRAF* mutational status such as mucinous histology and/or poor differentiation.^41^ However, we also observed that the highly scoring patches identified by the TCGA model failed to represent classical histopathological features of *BRAF* mutation and indeed 7 out of 9 highly scoring tiles in this group showed no tumor tissue or abundant artifacts (**Figure 4C**). The observation that such low-information patches were flagged by the model as being highly relevant shows that a model trained on TCGA only does not adequately learn to detect relevant patterns. Conversely, both swarm learning models identified plausible histopathological patches (i.e. patches with an expected phenotype) in 9 out of 9 highly scoring patches for *BRAF* mutation. Together, these data show that swarm learning-based AI models can generate predictions which are explainable and plausible to human experts even if not all of the single-cohort models trained on the same patient cohorts exhibit this behavior.

**Figure 4:**
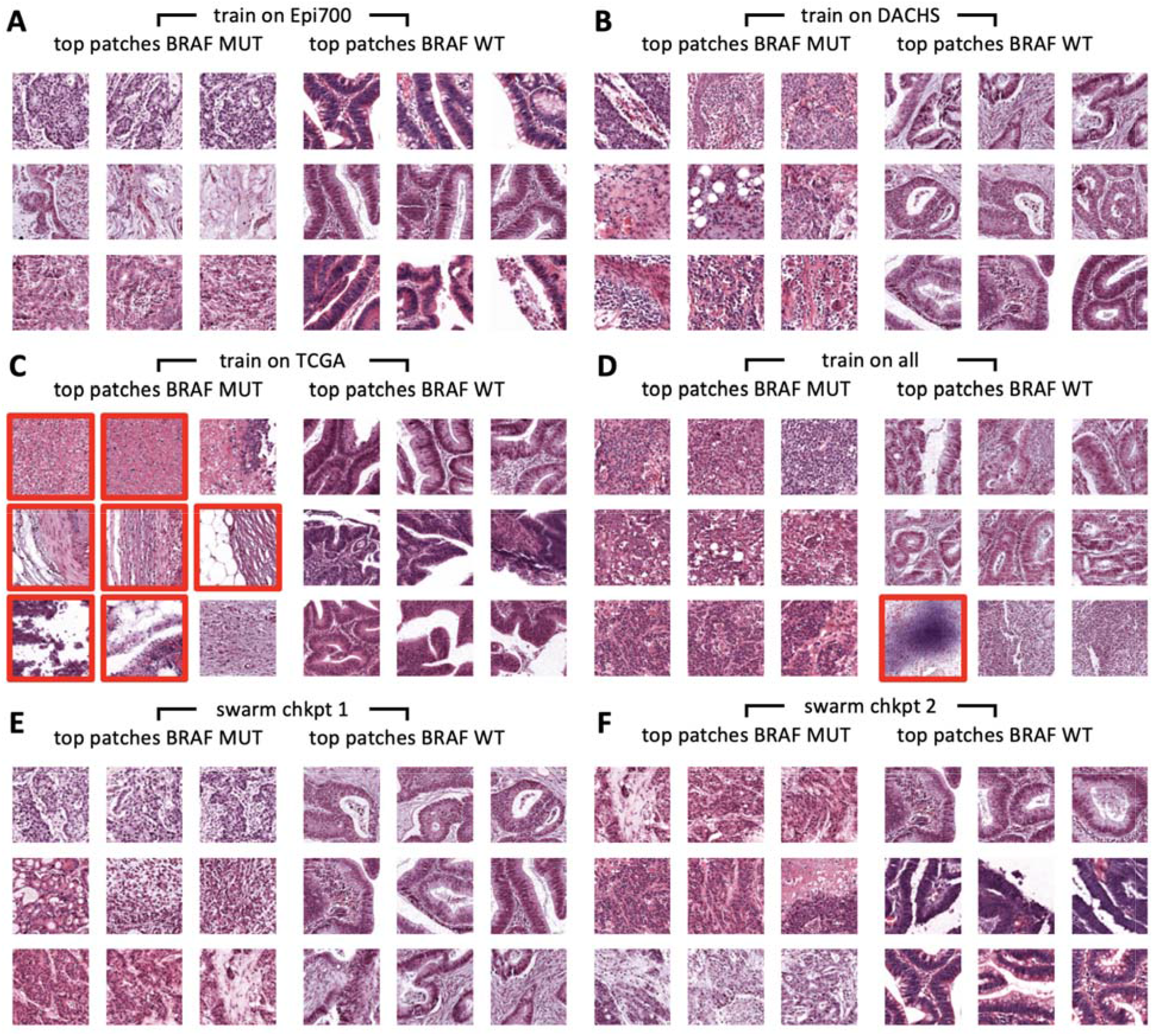
Highly predictive image patches for BRAF prediction. All patches were derived from QUASAR and were selected by the median model of five models which were trained on N=300 randomly selected patients per training cohort. (A) Model trained on Epi700, (B) trained on DACHS, (C) trained on TCGA, (D) trained on all three datasets, (E) swarm chkpt 1, (F) swarm chkpt 2). Red borders highlight tiles with artifacts or more than 50% non-tumor tissue.

## Discussion

Currently, the total amount of healthcare data is increasing at an exponential pace. In particular in histopathology, institutions across the world are digitizing their workflows, generating an abundance of data.^6^ These image data can be used in new ways, for example to make prognostic and predictive forecasts, aiming to improve patient outcomes.^3^ However, AI requires large and diverse datasets and its performance scales with the amount of training data.^19,20^ To train useful and generalizable AI models, institutions should be able to collaborate without jeopardizing patient privacy and information governance. In 2016, federated learning (FL) was proposed as a technical solution for such privacy-preserving distributed AI.^42^ FL enables joint training of AI models by multiple partners who cannot share their data with each other. However, FL relies on a central coordinator who monopolizes the resulting AI model, concentrating the power of exploitation in the hands of a single monopolistic entity. Thus, FL removes the need for data sharing but does not solve the problem of information governance. Swarm learning (SL), however, offers a solution to the governance problem, providing a true collaborative and democratic approach in which partners communicate and work on the same level, jointly and equally training models and sharing the benefits.^25,26,43^ Most recently, SL has been tested to detect COVID-19, tuberculosis, leukaemia and lung pathologies from transcriptome analysis or x-ray images, respectively^26^. In the present study, we demonstrate for the first time that the use of SL can enable AI-based detection of clinical biomarkers in solid tumors and yields high-performing models for pathology-based prediction of *BRAF* and MSI status, two important prognostic and predictive biomarkers in CRC.^3,9,44^

A possible technical limitation of our study is that we did not explicitly investigate differential privacy, although this could be incorporated in future work. Although histological images without their associated metadata are not considered protected health information (PHI) even under the Health Insurance Portability and Accountability Act (HIPAA) in the United States^45^, any membership inference attack or reconstruction of original data from shared model weight updates can be precluded by implementing additional differential privacy measures.^46^ Another limitation of this work is that the model performance needs to be further improved before clinical implementation. Previous work has shown that by increasing the sample size to ~10000 patients, classifier performance will increase.^19,20^ Our study shows that SL enables multiple partners to jointly train models without sharing data, thereby potentially facilitating the collection of such large training cohorts. Finally, previous proof-of-concept studies on SL in medical AI relied on virtual machines on a single bare-metal device. In the present study, we improved this by using three physically separate devices and implementing our code largely with open source software. While this indicates that SL is feasible between physically distinct locations, embedding SL servers in existing healthcare infrastructure in different institutions in multiple countries would probably require substantial practical efforts which should ideally be addressed in research consortia. To assess interchangeability of model data generated by swarm learning projects, validation of this technology in large-scale international collaborative efforts is needed. Our study provides a benchmark and a clear guideline for such future efforts, ultimately paving the way to establish SL in routine workflows.

## Additional information

### Author contributions

OLS, DT and JNK designed the study; OLS, NGL and JNK developed the software; OLS and TS performed the experiments; OLS and JNK analyzed the data; OLS, DC and NGL performed statistical analyses; PQ, MBL, MST, TJB, HIG, GGAH, EA, JAJ, RG, JCC, HB, MH and NPW provided clinical and histopathological data; all authors provided clinical expertise and contributed to the interpretation of the results. AS, TL, MH and CT provided resources and supervision. OLS, HSM and JNK wrote the manuscript and all authors corrected the manuscript and collectively made the decision to submit for publication.

## Acknowledgements

The authors are grateful to the customer support team of Hewlett Packard Enterprise (HPE) for providing support in using the HPE Swarm Learning package.

