## Supplementary material for "Swarm learning for decentralized artificial intelligence in cancer histopathology": Suppl. Data

### Supplementary Figures

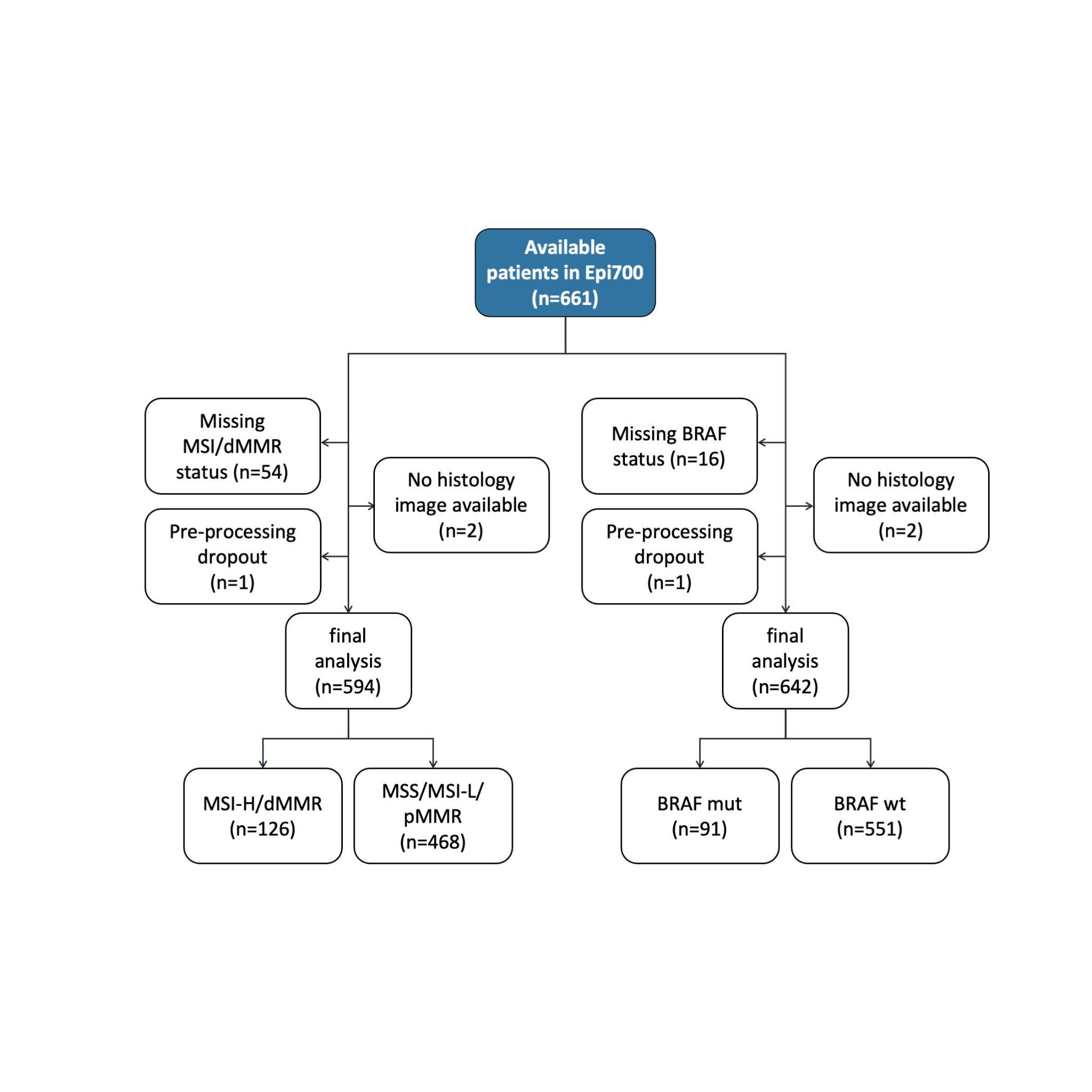

**Suppl. Figure S1: CONSORT chart for Epi700.**

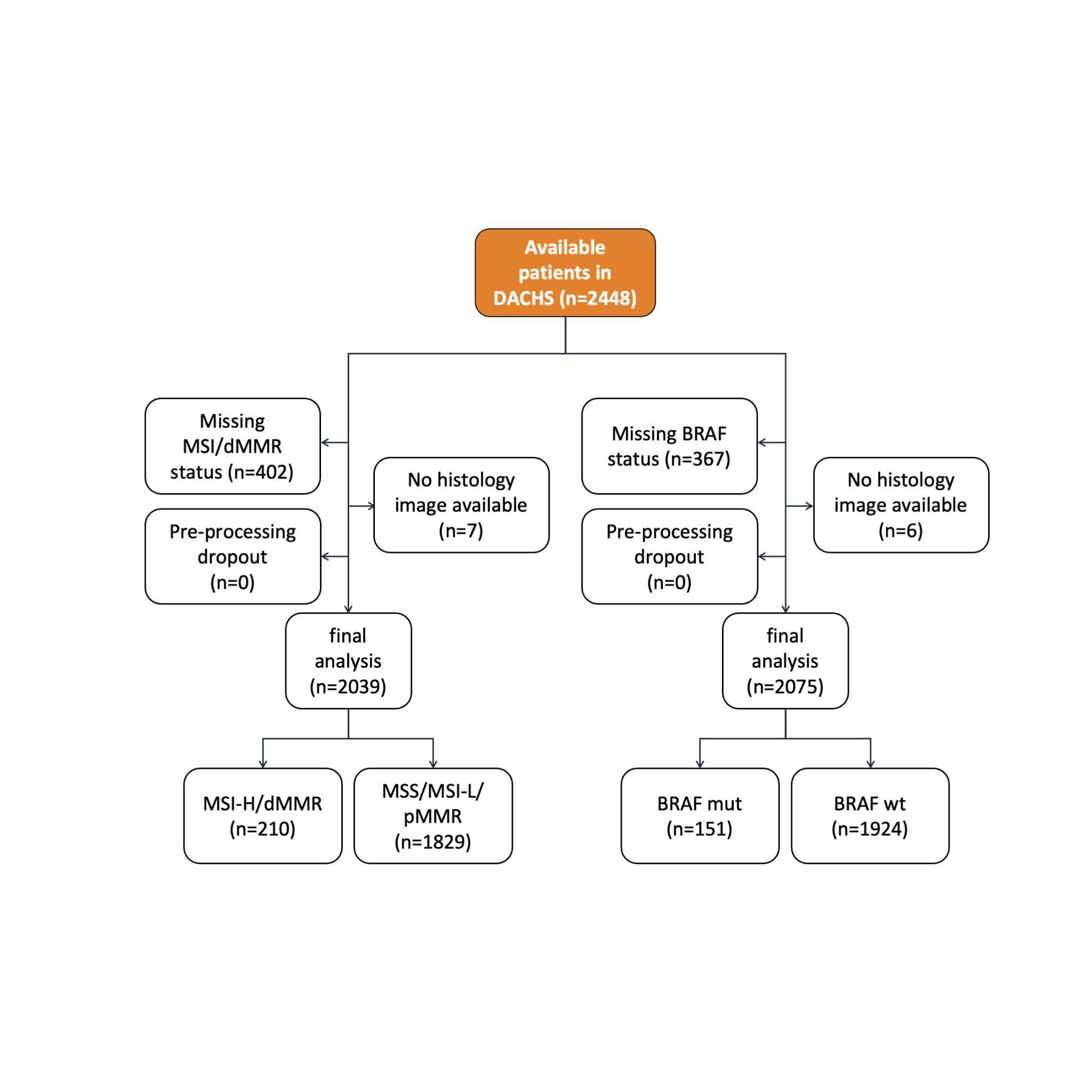

**Suppl. Figure S2: CONSORT chart for DACHS.**

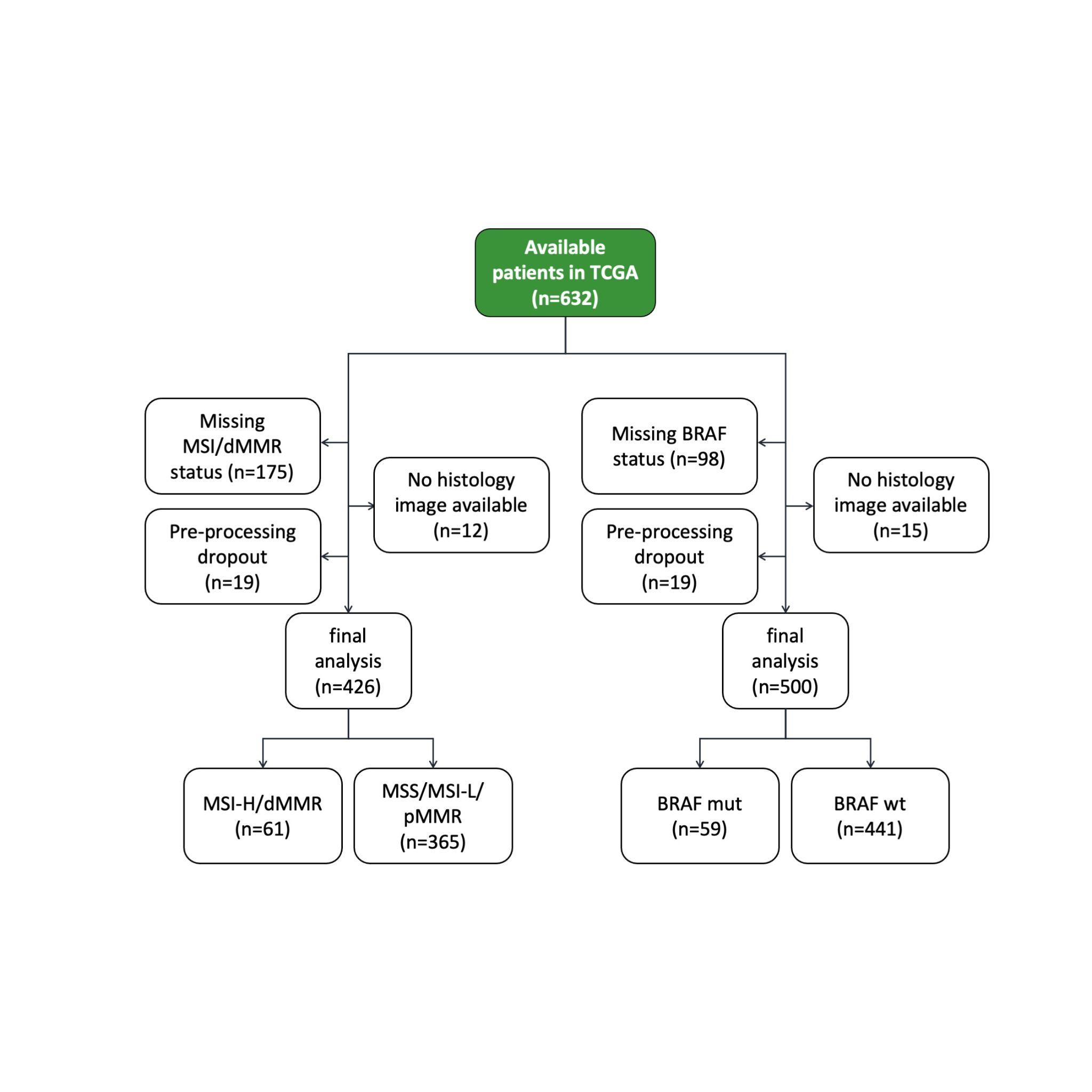

**Suppl. Figure S3: CONSORT chart for TCGA.**

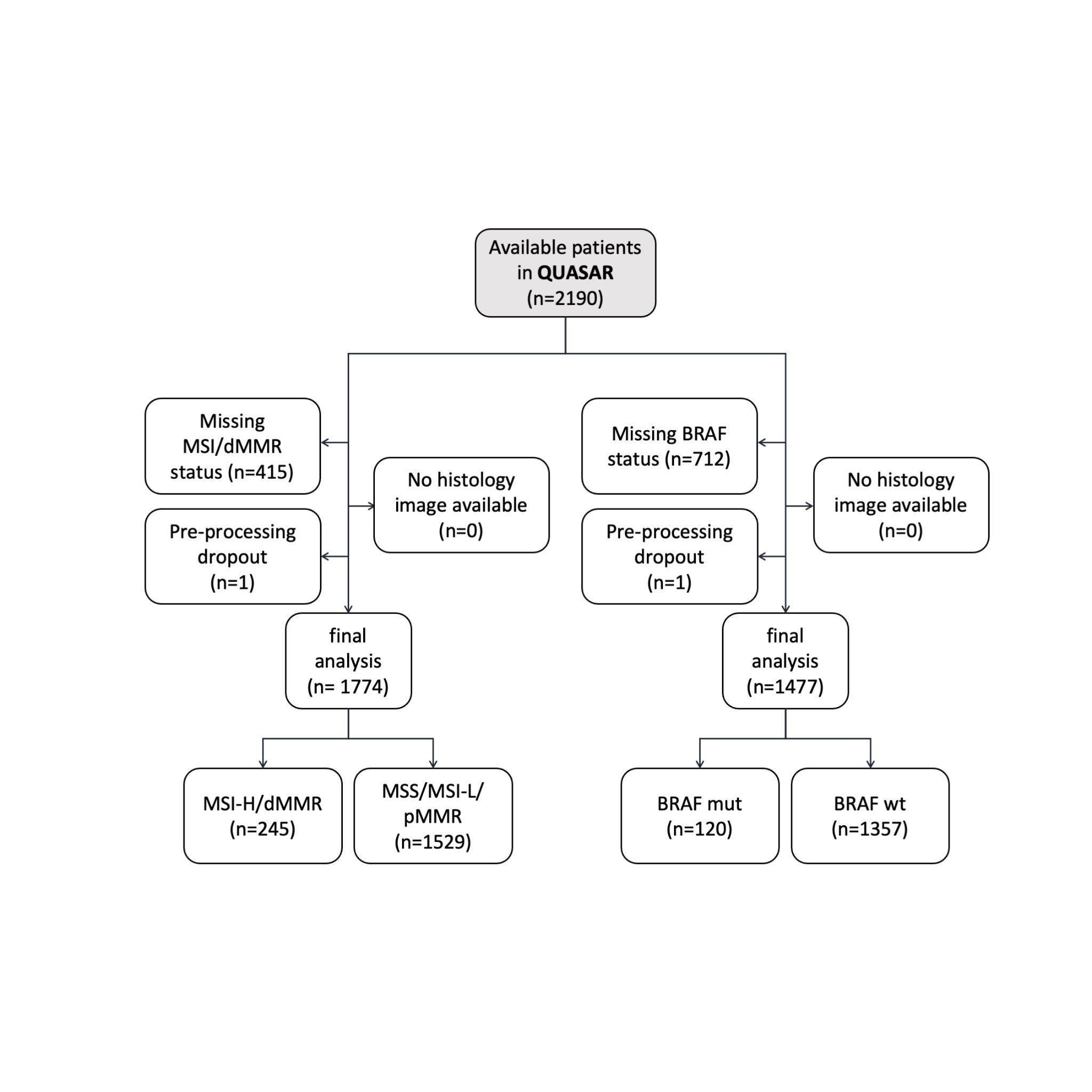

**Suppl. Figure S4: CONSORT chart for QUASAR.**

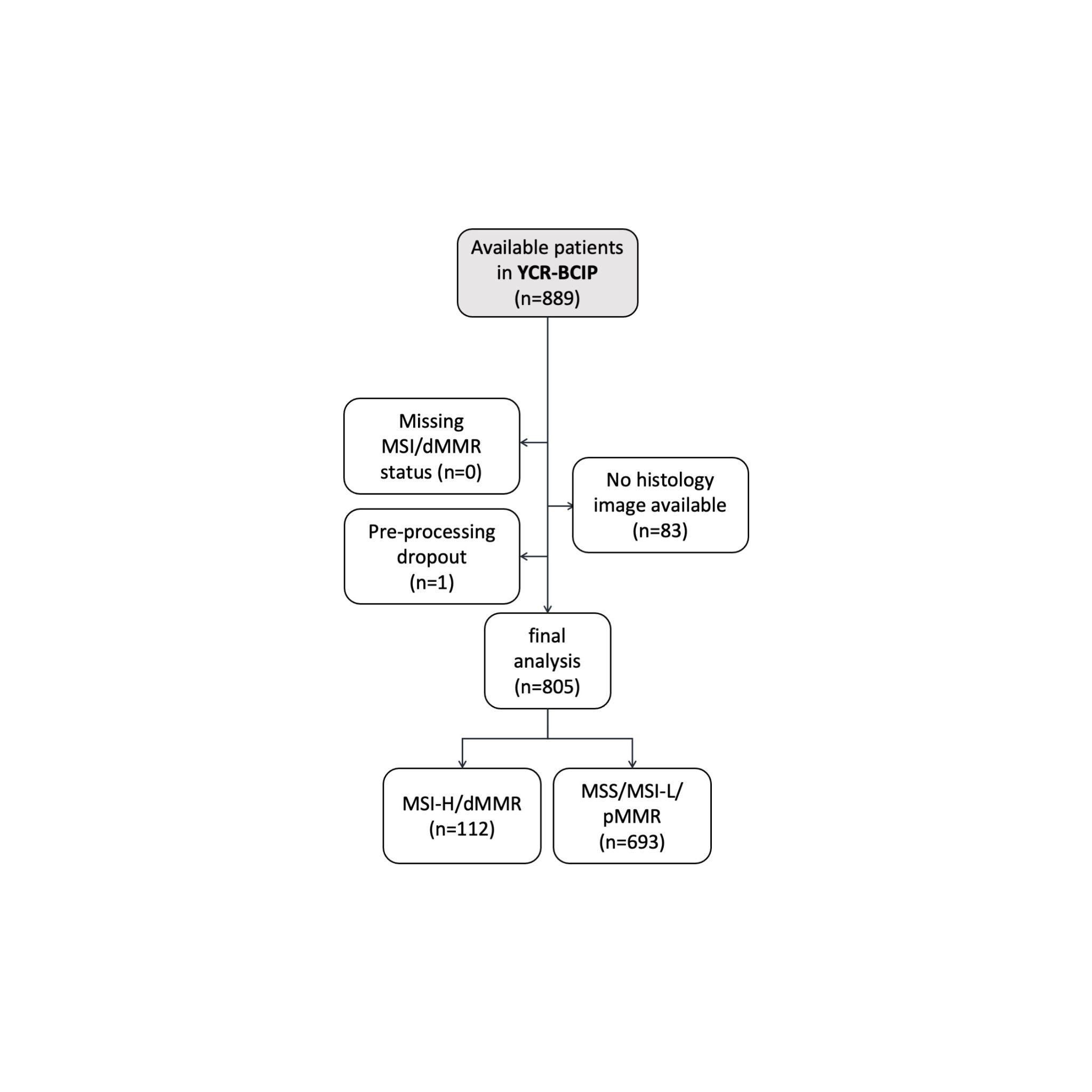

**Suppl. Figure S5: CONSORT chart for YCR-BCIP.**

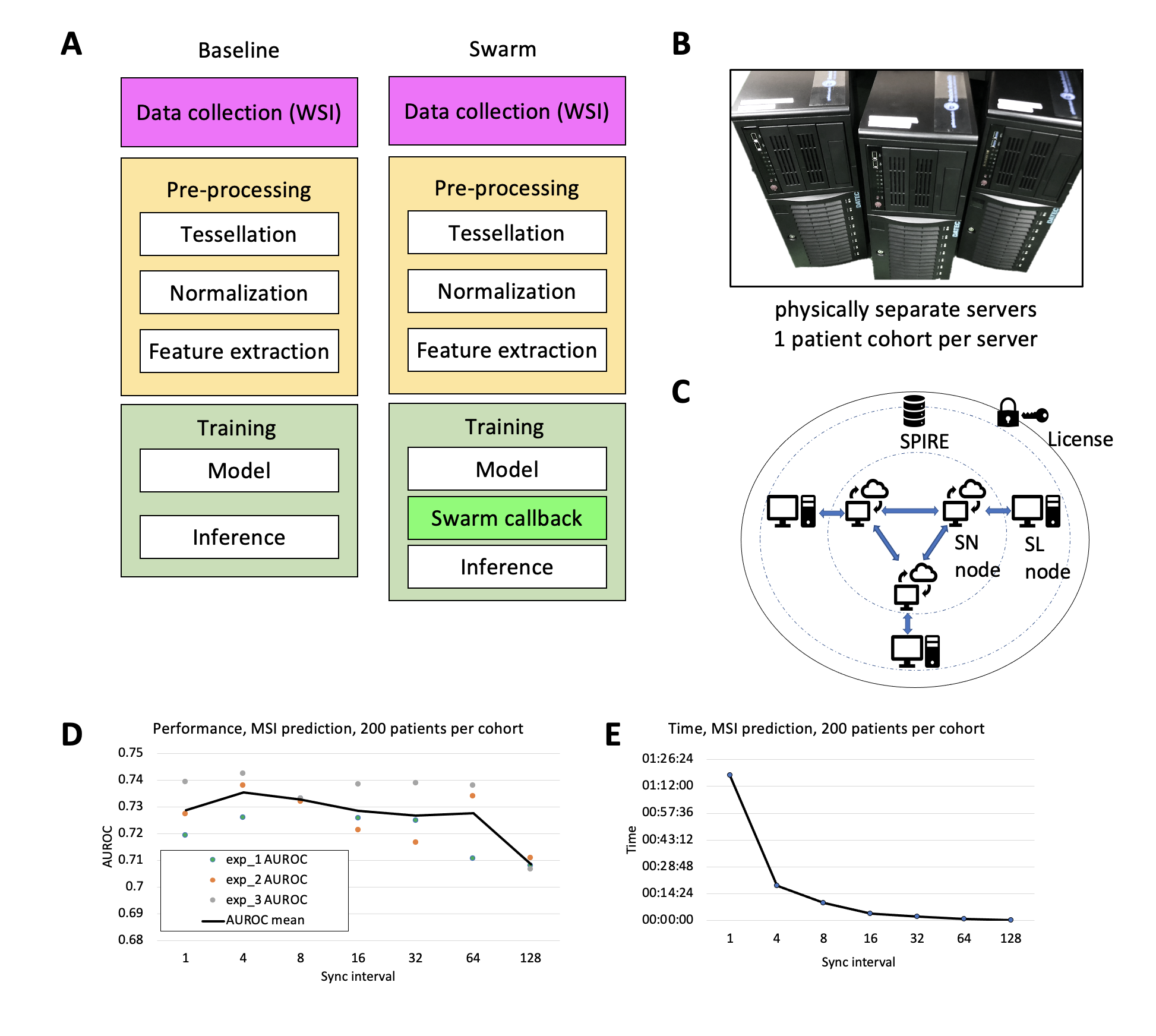

**Suppl. Figure S6: Workflow details and hyperparameter optimization. (**A) Schematic of baseline and SL-enabled workflow, (B) Physical server setup, (C) Schematic of HPE based SL^26^. (D) Hyperparameter optimization for sync interval based on the performance, (E) Time for training with different sync internal. Abbreviations: WSI = whole slide images, MSI = microsatellite instability, SL = swarm learning, SN = swarm network, SPIRE = SPIFFE Runtime Environment.

### Supplementary Tables

| **Category** | **Hyperparameter** | **Value** | **Reference** |
| --- | --- | --- | --- |
| HIA | Learning Rate | 0.0001 | Ghaffari Laleh et al.^37^ |
| HIA | Batch size | 124 |  |
| HIA | Number of Epochs | 5 |  |
| HIA | Optimizer | Adam |  |
| HIA | Network model | ResNet18 |  |
| Swarm | Sync interval | 4 | hyperparameter optimization |
| Swarm | min peers | 2 | https://github.com/HewlettPackard/swarm-learning |
| Swarm | adaptive sync | false |  |
| Swarm | node weights | 1 |  |

**Suppl. Table S1: HIA and swarm learning hyperparameters.**

[separate XLSX file]

**Suppl. Table S2. All performance results of all experiments, related to Figure 2 and Figure 3.**

|  | **Epi700** | **DACHS** | **TCGA** | **Merged** | **b-chkpt1** | **b-chkpt2** | **w-chkpt** |
| --- | --- | --- | --- | --- | --- | --- | --- |
| **Epi700** | 1 | 0.735447 | 0.054133 | 0.072777 | 0.008285 | 0.010536 | 0.001469 |
| **DACHS** | 0.735447 | 1 | 0.056813 | 0.019865 | 0.000511 | 0.000663 | 8.65E-05 |
| **TCGA** | 0.054133 | 0.056813 | 1 | 0.004342 | 0.000997 | 0.001145 | 0.000358 |
| **Merged** | 0.072777 | 0.019865 | 0.004342 | 1 | 0.343359 | 0.439399 | 0.037475 |
| **b-chkpt1** | 0.008285 | 0.000511 | 0.000997 | 0.343359 | 1 | 0.673565 | 0.015439 |
| **b-chkpt2** | 0.010536 | 0.000663 | 0.001145 | 0.439399 | 0.673565 | 1 | 0.00812 |
| **w-chkpt** | 0.001469 | 8.65E-05 | 0.000358 | 0.037475 | 0.015439 | 0.00812 | 1 |

**Suppl. Table S3: P-values for pairwise comparisons between experiments for BRAF prediction in QUASAR for all patients** (unpaired, two-sided t-test).

|  | **Epi700** | **DACHS** | **TCGA** | **Merged** | **b-chkpt1** | **b-chkpt2** | **w-chkpt** |
| --- | --- | --- | --- | --- | --- | --- | --- |
| **Epi700** | 1 | 0.300247 | 1.54E-05 | 0.001257 | 0.015433 | 8.79E-05 | 8.93E-06 |
| **DACHS** | 0.300247 | 1 | 0.64237 | 0.064677 | 0.199045 | 0.112427 | 0.057049 |
| **TCGA** | 1.54E-05 | 0.64237 | 1 | 5.80E-05 | 1.09E-05 | 6.14E-07 | 2.83E-07 |
| **Merged** | 0.001257 | 0.064677 | 5.80E-05 | 1 | 0.009638 | 0.120348 | 0.864977 |
| **b-chkpt1** | 0.015433 | 0.199045 | 1.09E-05 | 0.009638 | 1 | 0.010362 | 0.000234 |
| **b-chkpt2** | 8.79E-05 | 0.112427 | 6.14E-07 | 0.120348 | 0.010362 | 1 | 0.008928 |
| **w-chkpt** | 8.93E-06 | 0.057049 | 2.83E-07 | 0.864977 | 0.000234 | 0.008928 | 1 |

**Suppl. Table S4: P-values for pairwise comparisons between experiments for MSI prediction in QUASAR for all patients** (unpaired, two-sided t-test).

|  | **Epi700** | **DACHS** | **TCGA** | **Merged** | **b-chkpt1** | **b-chkpt2** | **w-chkpt** |
| --- | --- | --- | --- | --- | --- | --- | --- |
| **Epi700** | 1 | 0.810466 | 0.562843 | 8.88E-05 | 0.06603 | 0.005866 | 4.33E-05 |
| **DACHS** | 0.810466 | 1 | 0.919589 | 0.169501 | 0.833261 | 0.575862 | 0.203239 |
| **TCGA** | 0.562843 | 0.919589 | 1 | 8.08E-05 | 0.135212 | 0.008523 | 2.87E-05 |
| **Merged** | 8.88E-05 | 0.169501 | 8.08E-05 | 1 | 0.000171 | 0.001703 | 0.464731 |
| **b-chkpt1** | 0.06603 | 0.833261 | 0.135212 | 0.000171 | 1 | 0.065841 | 4.10E-05 |
| **b-chkpt2** | 0.005866 | 0.575862 | 0.008523 | 0.001703 | 0.065841 | 1 | 0.000734 |
| **w-chkpt** | 4.33E-05 | 0.203239 | 2.87E-05 | 0.464731 | 4.10E-05 | 0.000734 | 1 |

**Suppl. Table S5: P-values for pairwise comparisons between experiments for dMMR prediction in YCR-BCIP for all patients** (unpaired, two-sided t-test).

|  | **Epi700** | **DACHS** | **TCGA** | **Merged** | **b-chkpt1** | **b-chkpt2** | **w-chkpt** |
| --- | --- | --- | --- | --- | --- | --- | --- |
| **Epi700** | 1 | 0.124088 | 0.15321 | 0.164606 | 0.578819 | 0.42984 | 0.208304 |
| **DACHS** | 0.124088 | 1 | 0.655429 | 0.000639 | 0.02226 | 0.008388 | 0.000813 |
| **TCGA** | 0.15321 | 0.655429 | 1 | 0.013389 | 0.059135 | 0.038283 | 0.016248 |
| **Merged** | 0.164606 | 0.000639 | 0.013389 | 1 | 0.339588 | 0.438796 | 0.778013 |
| **b-chkpt1** | 0.578819 | 0.02226 | 0.059135 | 0.339588 | 1 | 0.807427 | 0.435173 |
| **b-chkpt2** | 0.42984 | 0.008388 | 0.038283 | 0.438796 | 0.807427 | 1 | 0.567631 |
| **w-chkpt** | 0.208304 | 0.000813 | 0.016248 | 0.778013 | 0.435173 | 0.567631 | 1 |

**Suppl. Table S6: P-values for pairwise comparisons between experiments for BRAF prediction in QUASAR for 200 patients** (unpaired, two-sided t-test).

|  | **Epi700** | **DACHS** | **TCGA** | **Merged** | **b-chkpt1** | **b-chkpt2** | **w-chkpt** |
| --- | --- | --- | --- | --- | --- | --- | --- |
| **Epi700** | 1 | 0.113646 | 0.381066 | 1.44E-05 | 0.021346 | 0.01977 | 0.008786 |
| **DACHS** | 0.113646 | 1 | 0.05067 | 3.55E-05 | 0.004533 | 0.004592 | 0.002457 |
| **TCGA** | 0.381066 | 0.05067 | 1 | 0.000443 | 0.274513 | 0.114665 | 0.059786 |
| **Merged** | 1.44E-05 | 3.55E-05 | 0.000443 | 1 | 7.55E-05 | 0.017031 | 0.029182 |
| **b-chkpt1** | 0.021346 | 0.004533 | 0.274513 | 7.55E-05 | 1 | 0.314582 | 0.152971 |
| **b-chkpt2** | 0.01977 | 0.004592 | 0.114665 | 0.017031 | 0.314582 | 1 | 0.730893 |
| **w-chkpt** | 0.008786 | 0.002457 | 0.059786 | 0.029182 | 0.152971 | 0.730893 | 1 |

**Suppl. Table S7: P-values for pairwise comparisons between experiments for MSI prediction in QUASAR for 200 patients** (unpaired, two-sided t-test).

|  | **Epi700** | **DACHS** | **TCGA** | **Merged** | **b-chkpt1** | **b-chkpt2** | **w-chkpt** |
| --- | --- | --- | --- | --- | --- | --- | --- |
| **Epi700** | 1 | 0.605611 | 0.072281 | 4.92E-05 | 0.229145 | 0.022538 | 0.005563 |
| **DACHS** | 0.605611 | 1 | 0.06765 | 0.000356 | 0.175667 | 0.0306 | 0.01444 |
| **TCGA** | 0.072281 | 0.06765 | 1 | 0.00249 | 0.302962 | 0.679152 | 0.458176 |
| **Merged** | 4.92E-05 | 0.000356 | 0.00249 | 1 | 6.96E-05 | 0.001888 | 0.00042 |
| **b-chkpt1** | 0.229145 | 0.175667 | 0.302962 | 6.96E-05 | 1 | 0.108632 | 0.026695 |
| **b-chkpt2** | 0.022538 | 0.0306 | 0.679152 | 0.001888 | 0.108632 | 1 | 0.746109 |
| **w-chkpt** | 0.005563 | 0.01444 | 0.458176 | 0.00042 | 0.026695 | 0.746109 | 1 |

**Suppl. Table S8: P-values for pairwise comparisons between experiments for dMMR prediction in YCR-BCIP for 200 patients** (unpaired, two-sided t-test).

### Supplementary Methods

#### Practical details on the swarm learning setup

For usage of the HPE swarm learning community edition, the initial step is to register an HPE license to run the SL platform from (https://myenterpriselicense.hpe.com/cwp-ui/auth/login). This process is managed by the Licence Server node. Docker containers are one of the important parts in the SL approach. There is a docker container image for all of the SL components which is pulled from the registry (https://github.com/HewlettPackard/swarm-learning/blob/master/docs/setup.md# pull-docker-images). A separate docker image is built on top of the existing docker image for the SL node with all the Python modules necessary for our workflow.

#### Swarm learning implementation

The principle of SL is to jointly train an AI model in multiple systems. Parameters are sent from each partner to the other peers at multiple sync stops, and that these are averaged at each sync, before the process moves on with all sources using the averaged parameters. We assume that there are three physically separate systems (System A, System B, System C) used for the SL framework in our approach. All images were available in ScanScope Virtual Slide (SVS) format and were tessellated using https://github.com/KatherLab/preProcessing according to the “The Aachen Protocol for Deep Learning Histopathology: A hands-on guide for data preprocessing”.^47^ The baseline Deep Learning prediction workflow was described previously.^37^ The training cohorts TCGA, Epi700 and DACHS trained on System A, System B and System C respectively. The training on the individual systems performed. 512 features from the resnet 18 model pre-trained on imagenet for the patches and its respective labels for the target (MSI/dMMR and BRAF) are cached and re-used as an input for the FCN. The framework (Pytorch) and model (HIA) remains the same in all the three systems. The SL process begins with the enrolment of nodes with the swarm network^26^, System A is used to initialize the licence server by starting the licence container and installing the swarm licence downloaded from the HPE login website. System A also starts the SPIFFE SPIRE container. The first SN process (node) to go online is referred to as the “sentinel” node and will be the first to register itself with the SPIFFE SPIRE network. When the SN node of the system A is ready the, SN node of System B and System C are run. During training each system trains its data batch in the local system till the merging criterion (sync interval) is reached. The node which finishes its training batch first will be the leader and will collect the learning from other peers (depending on the minimum number of peers, in our case two), average the learning weights and send it back. Since the data samples are of different sizes the SL node stops at different instances creating different checkpoint models.

#### Hardware

In our setup, the three systems used for training individual models and SL had the following hardware setup: System A with a 128 GB RAM, two GPUs NVIDIA Quadro RTX 6000 with 24 GB RAM. System B with 64 GB RAM, one GPU NVIDIA RTX A6000 with 48 GB RAM. System C with 64 GB RAM, two GPUs NVIDIA Quadro RTX 6000 with 24 GB RAM All the three systems connected to a 1 GBit/sec Ethernet port with three open ports for communication, ran Ubuntu 20.04, Docker container, passwordless SSH and synchronized time across all systems.

#### Hyperparameter optimization

The sync interval describes how many training iterations elapse before all partners exchange and average the model weights. The following values were explored: 1, 4, 8, 16, 32, 64 and 128 with training time and AUROC values as primary endpoints. For training of the models used in hyperparameter optimization, 200 patients per cohort were used from Epi700, DACHS and TCGA each and performance was assessed on the QUASAR and YCR-BCIP cohort.
